# Optimized sgRNA design by deep learning to balance the off-target effects and on-target activity of CRISPR/Cas9

**DOI:** 10.1101/2020.03.04.976340

**Authors:** Jie Lan, Yang Cui, Xiaowen Wang, Guangtao Song, Jizhong Lou

**Author notes:** To whom correspondence should be addressed. Correspondence may also be addressed to Guangtao Song.

## Abstract

The CRISPR/Cas9 system derived from bacteria especially *Streptococcus pyogenes* (*Spy*Cas9) is currently considered as the most advanced tool used for numerous areas of biological study in which it is useful to target or modify specific DNA sequences. However, low on-target cleavage efficiency and off-target effects impede its wide application. Several different sgRNA design tools for *Spy*Cas9 by using various algorithms have been developed, including linear regression model, support vector machine (SVM) model and convolutional neuron network model. While the deep insight into the sgRNA features contributing for both on-target activity and off-target still remains to be determined. Here, with public large-scale CRISPR screen data, we evaluated contribution of different features influence sgRNA activity and off-target effects, and developed models for sgRNA off-target evaluation and on-target activity prediction. In addition, we combined both activity and off-target prediction models and packaged them as an online sgRNA design tool, OPT-sgRNA.

## INTRODUCTION

The clustered regularly interspaced short palindromic repeats (CRISPR) genome engineering technology has emerged as a powerful molecular tool for genome editing with promising therapeutic and research advancements (1–7). Among the many Crispr and their associated (Crispr-cas) protein systems, Cas9 and Cas12a are adopted to use for genome editing. As a single protein adaptor, they use small noncoding RNAs as guide to recognize and edit the target sites. The molecular mechanism of DNA recognition and cleavage of these systems have been studied in-depth (8–16). Cas9 from *Streptococcus pyogenes* (SpyCas9, referred as Cas9 hereafter) are the most widely used. By simply changing the sequence of the single-guide RNA (sgRNA), we can easily reprogram Cas9 to target different sites in the genome. However, various studies showed that CRISPR/Cas9 displayed a wide variety of activities across different target sites, leading to the conclusion that some target sites are inherently more effective (17–20). Therefore, screening of sgRNAs with maximum on-target activity and minimum potential off-target effects is a glorious but challenging task.

Recently, more than 26 computational approaches have been developed for sgRNA design and evaluation (20–24). These tools are designed to assist researchers in the selection of best target sites by helping them exclude undesirable targets based on predicted low efficiency or a high potential for off-target effects. They could be broadly divided into two groups, on-target cleavage efficiency tools (6,25–39) and off-target activity tools (18,28,33,40–51). In the on-target cleavage efficiency evaluation tools, researchers focus on identifying the gRNA sequence features that contribute to target cleavage efficiency. Despite the differences in their mathematical model, some common key rules have been identified. These include a G preceding the PAM, avoiding poly-T sequences and limiting the GC content (39,52–54). In addition, a PAM-proximal seed region has been established to be critical for Cas9 activity (1,54). More recent studies also pay attention to non-sequence information, such as location of the sgRNA target site within the gene and the thermodynamic stability of the sgRNA (39,45,55,56). In the off-target effect prediction tools, people usually first align the target site to the reference genome, and identify similar locations that may be inadvertently targeted by Cas9. These potential off-target sites were then evaluated by experimental studies, and to train scoring algorithms, which attempt to predict the potential off-target activity. The two most popular scoring algorithms are MIT Broad score (18) and CFD score (33). Both scoring methods are based on evaluating the contributions made by different mismatch positions and number in the target site, and calculating a weight matrix to determine off-target efficiency of each sgRNA.

Although much progress has been made for the sgRNA design and evaluation, very few tools which could implement both on-target activity and off-target effect evaluation are available. Here, we present a new computational tool to combine off-target prediction and on-target activity evaluation. We incorporate new computational design rules in both off-target and on-target scoring algorithm. In addition, we create a web-based tool OPT-sgRNA to enable optimized sgRNA design that maximize on-target activity and minimize off-target effects.

## METHODS

### Data assembly

The training dataset for sgRNA off-target evaluation was assembled from published data obtained by Keith Joung’s group and Jennifer Doudna’s group using GUIDE-seq methods (57–59). The data in their experiments are composed of 19 sgRNAs and 753 off-target site pairs in total. The training dataset for sgRNA on-target activity effect was downloaded from GenomeCRISPR database and chose only one dataset considering batch effect among different source (60). In this dataset, Aguirre and colleagues performed genome-scale loss-of-function screens in 33 cancer cell lines to identify genes essential for proliferation/survival (61). From that analysis, essential genes have been discovered. To avoid bias in different cell lines, we selected the sgRNA expressing data of A375 cell line with 118,862 sgRNAs targeting 22,329 genes. In recent studies, researchers found that sgRNA with polyT end with premature of RNA and low GC-content end with lower activity. Therefore, sgRNAs with polyT (TTTT) and GC-content lower than 40% are excluded. In addition, we chose the most potent sgRNAs (top 10% in ranking) and the least potent sgRNAs (bottom 10%). In totally, 40,234 sgRNAs targeting 19,561 genes were used for the training.

### Potential sgRNA scanning and corresponding Off-target site searching

For a given genes, we first scan its exon and promoter regions with NGG site. SeqMap (62) is a tool for mapping large amount of short sequences to genome and designed to find all the places in the reference genome where sequence may come from. Each of sgRNA is aligned to genome using SeqMap (v 1.0.12) to detect its off-target site with no more than 4 mismatches.

### sgRNA activity and off-target prediction model

With a large number of features (1538), we incorporate an L1-regularized linear support vector machine (SVM) to generate subsets of features with the best generalization error. With different parameters applied in feature selection (0.01, 0.05, 1), increased features help little to increase final model accuracy. Given the subset of features from the feature selection by L1-SVM (240 of 1538), we implemented several model (Gradient Boosting, Random Forest, Logistic Regression, Decision Tree, KNeighbors, Neural Net, Naïve Bayes, linear SVM and RBF SVM) and evaluated their performance by 10-fold cross validation. All of the sgRNAs were randomly into 10 parts and then nine parts were set as training dataset and the remained part is testing dataset. By 10 rounds of training and testing process, all of the accuracy of models were calculated by mean of each round. With the AUC (area under curve) of the ROC curve, Logistic Regression performed as best model and then applied to calculate weights for each selected features. With the weight from Logistic Regression model, sgRNAs activity score can be easily computed. A sgRNA with a subset of all the features then transformed to binary variables by one-hot encoding. *S_j_* is sgRNA j and ω*_i_* is weight of feature i. The activity score of a given sgRNA j is provided via logistic regression as:

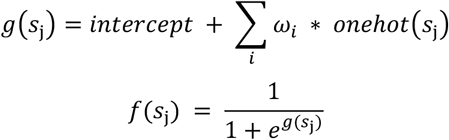

In the end, sgRNA activity score *f*(*s_j_*) will fall into range of 0 to 1, and higher values means higher activity.

It is known that mismatch is the most important feature decease the effect on off-target sites, we perform a linear regression model to detect effects of different regions (the number of mismatches in 1-2, 3-8, 9-15, 16-18 and 19-20 nucleotides ahead PAM). For a given sgRNA, we use the linear model to predict activity of sgRNAs’ off-target sites one by one and the whole off-target effect is measured by the sum of effect of each off-target site.

### Comprehensive library construction

Human (hg38) and Mouse (mm10) genomes and the corresponding gene annotation files are downloaded from NCBI and promoter sequences (upstream2000) are downloaded from UCSC genome database. For a given gene, we first scan sgRNAs in its exon and promoter regions for PAM sequence on both the forward and reverse strand, and then search off-target sites in the remaining regions using SeqMap with mismatches<=4. In all, the major process of sgRNAs evaluation can be divided into two parts: first we sort all of sgRNAs by their off-target score and choose bottom 100 as the low off-targets sgRNAs. Next, activity score is evaluated by logistic regression model and select only top 10 sgRNAs measured by their activity scores. All of the available library can be downloaded from our webserver OPT-sgRNA.

## RESULTS

### Flow of the sgRNA screening and evaluation procedures

To overcome the limitations of currently available sgRNA design tools, we designed a new online sgRNA predictor—OPT-sgRNA, suitable for editing for human or mouse genome using SpyCas9. As seen in Figure 1, for any provided gene name or DNA sequence, OPT-sgRNA first scans sequence of all exons by genomic coordinate and their corresponding promoters with PAM sequence. sgRNAs with polyT or low GC-content are filtered out before further evaluation. Next, for each sgRNA, all of its off-target sites are retrieved by aligning it to whole genome excluding regions of query gene by SeqMap (62) with mismatches setting as 4, and then the off-target effect is measured by the sum of all off-target site effect evaluated by linear regression model. All of sgRNAs are then sorted incrementally by its off-target effect and only top M sgRNAs (The default M is 100 and it can be set by user) with lower off-target effect are selected for the next activity scoring. sgRNAs scoring process are then started quickly and sgRNAs are ordered by their activity, top N sgRNAs with higher activity are presented as output (The default N is 10 and it can also be set by user).

**Figure 1.**
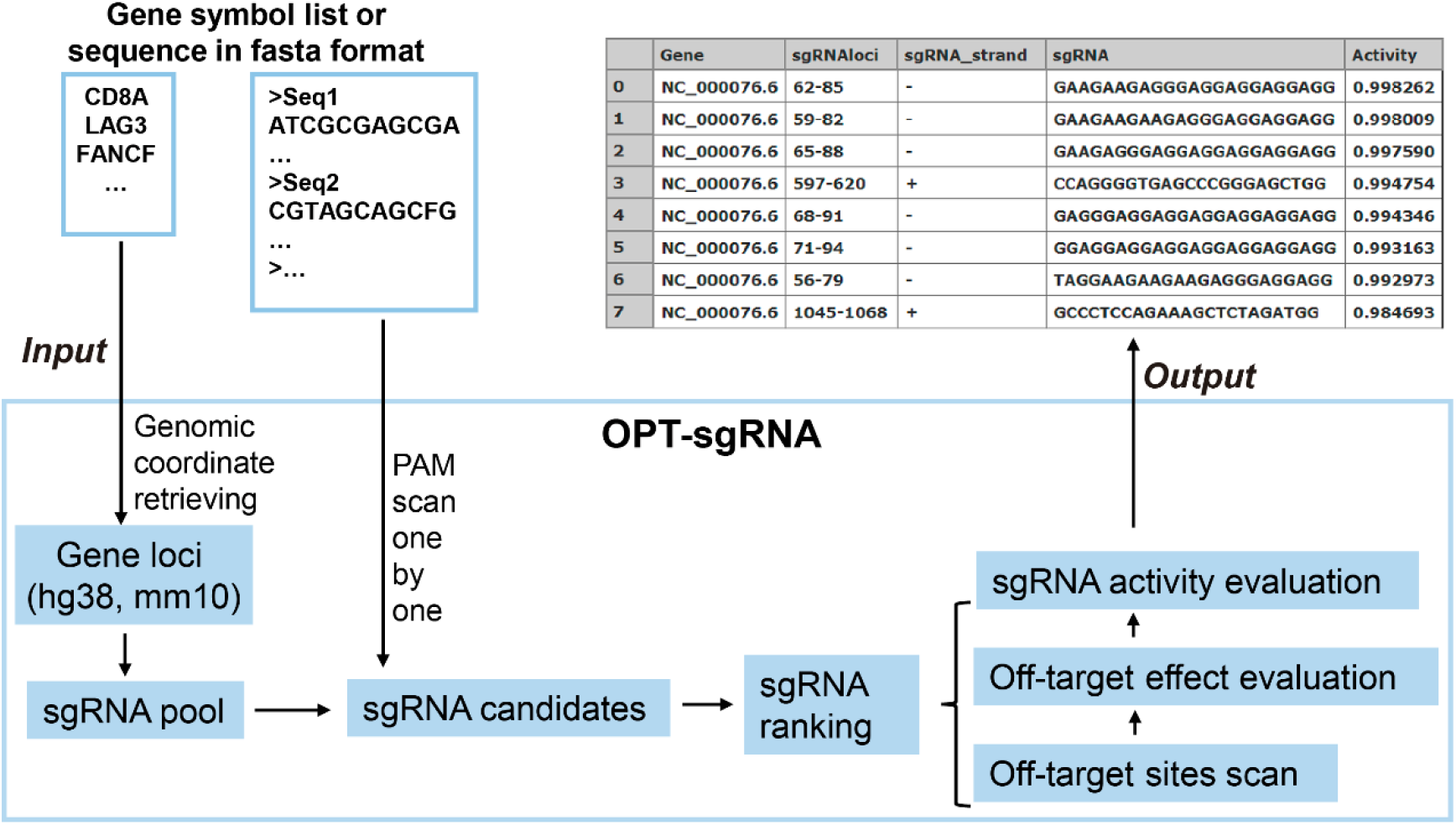
The scheme of OPT-sgRNA design tool. A list of gene symbol or sequence in fasta format could be used as the input for OPT-sgRNA. The number of genes or sequences is not limited. First, OPT-sgRNA performs PAM scanning for the given sequences or retrieves sgRNAs for the given genes in pre-constructed sgRNA pool(hg38, mm10). Second, OPT-sgRNA ranks all of the sgRNA candidates and then starts the evaluation. Off-target sites will be detected along genomes excepts target gene region with no more than 4 mismatches between off-target site and specified sgRNA. All of the off-target sites aggregated as an off-target score and sgRNAs sorted by off-target score in increasing manner, top M sgRNAs activity are evaluated and top N sgRNAs will be output to the interface (M and N are default by 100 and 10, respectively. Both M and N can be set by user).

### Evaluation of off-target effect

Several computational methods already exist to predict off-target sites and/or evaluate the specificity of the sgRNAs (18,28,33,41–50). Two main features are used to predict the specificity of the sgRNA: number and loci of mismatches, binding energy between sgRNA and target DNA. However, previous structural and biochemical studies have shown that the sgRNA-Cas9 complex could divide target DNA into several distinct regions: linker, seed, middle and tail region (8,10,12,15). Our previous biochemical studies showed that the mismatch between sgRNA and target DNA in these regions show very different effect on target cleavage efficiency (15). Therefore, previous simple number of mismatches feature is too crude for the accurate evaluation. Here we use the GUIDE-Seq dataset from Keith Joung group and Jennifer Doudna group to perform off-target effect training (58,59). The dataset contains 753 off-target sequences reported for 19 different gRNAs. We choose number of mismatches in 5 different regions as factors to training the off-target prediction model (Figure 2). It could be found that when the number of mismatches is higher than 4, the cleavage activity of Cas9 decreased significantly (Figure 2A). And as we expected, the position of mismatch is also a very important parameter to evaluate the specificity of sgRNAs. The seed region shows the most significant effect on target cleavage (Figure 2C). We next constructed a new Linear Regression model to calculate the off-target score of a specific off-target site (Figure 2D).

**Figure 2.**
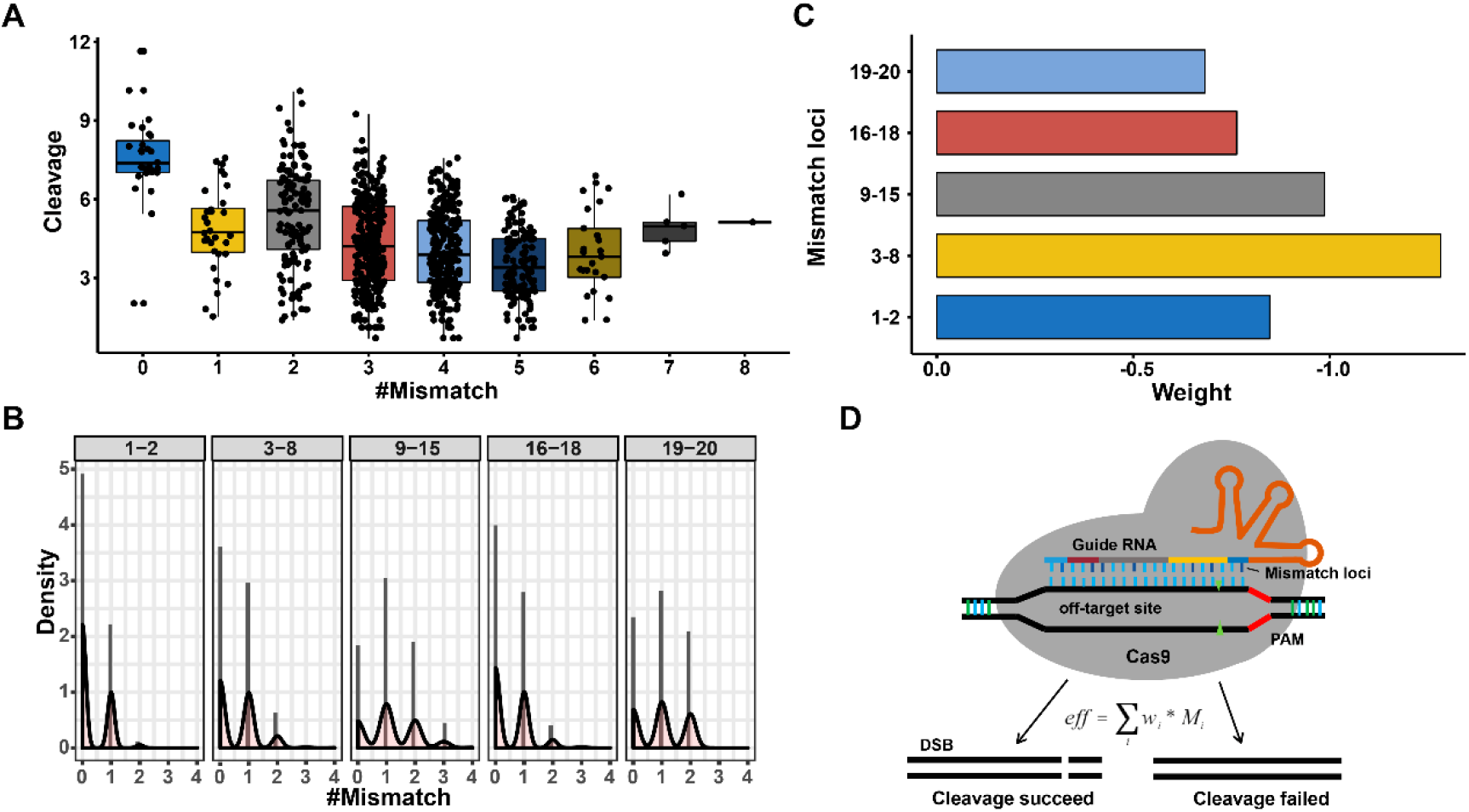
The impact of mismatch position on the off-target effect. **(A)** Cleavage efficiency of 19 sgRNAs on 753 off-target sites with different numbers of mismatches. The sgRNA and off-target site pair with 6 or more mismatches is rarely observed. **(B)** Number of mismatches in different regions. sgRNA can be separated into 5 different regions with discriminate effect to target activity and off-target specificity. The number of mismatches is shown in x-axis, y-axis is the observed density of number of mismatches. **(C)** The weight of different mismatch loci depicted with linear regression model. **(D)** The off-target evaluation model. For a given pair of sgRNA and off-target site, the number of mismatches in the specified regions is extracted and off-target activity will be calculated.

### sgRNA activity evaluation and selection

After classification of potential off-target sites, candidate sgRNAs with minimized off-target effects can be further evaluated by their activity. As seen in Figure 3, The sgRNA activity dataset contains 118,862 sgRNAs targeting 22,329 genes. SgRNAs with polyT (TTTT) and GC-content lower than 40% are excluded and only the most potent sgRNAs (top 10% in ranking) and the least potent sgRNAs (bottom 10%) are selected as finally sgRNA dataset. In all, there are 19561 genes with 40234 sgRNAs. Next, features such as single nucleotide, neighboring di-nucleotides and tri-nucleotides, GC-content are extracted and transformed by one-hot coding (Figure 3, right panel) for feature selection by L1-SVM. Finally, 10-folds cross validation performed for model selection. To build an efficient prediction model, we need to select important features and discard all irrelevant features. There are various existing machine learning methods such as wrapper or filter method that we can apply to do this job. With L1-SVM applied to feature selection, number of features deceases from 1538 (original) to 240 (finally). It’s interesting that accuracy does not increase with more features used (parameters set as 0.1, 0.05, 0.01) (Figure S1, Figure 4A). Considering different models applied in recent sgRNA selection tools, we train each model on selected features and evaluate their performance by 10-fold cross validation. By all, the logistic regression classifier performs as the best model (accuracy and robustness) to predict sgRNA activity accurately. To our surprise, in addition to single- or di-nucleotides preferences as reported before (Figure S2) (54,55). We also observed tri-nucleotides preference in our model (Figure 4B).

**Figure 3.**
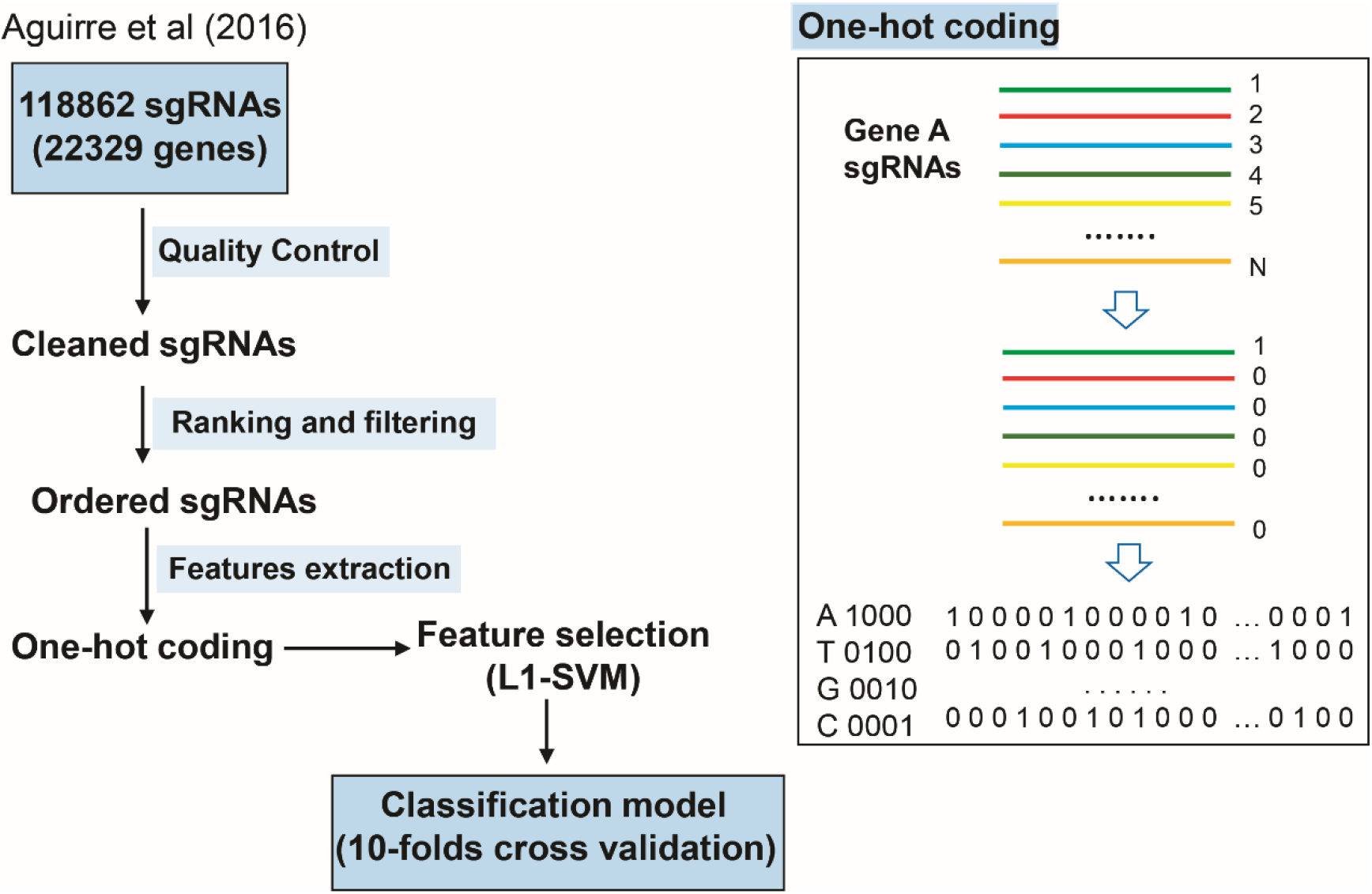
The training and evaluation workflow of the sgRNA activity model. The sgRNA activity dataset contains 118,862 sgRNAs targeting 22,329 genes. First, sgRNAs with polyT (TTTT) and GC-content lower than 40% are excluded and only the most potent sgRNAs (top 10% in ranking) and the least potent sgRNAs (bottom 10%) are selected as final sgRNA dataset. In all, there are 19561 genes with 40234 sgRNAs. Next, features such as single nucleotide, neighboring di-nucleotides and tri-nucleotides, GC-content are extracted and transformed by one-hot coding (right panel) for feature selection by L1-SVM. Finally, 10-folds cross validation performed for model selection.

**Figure 4.**
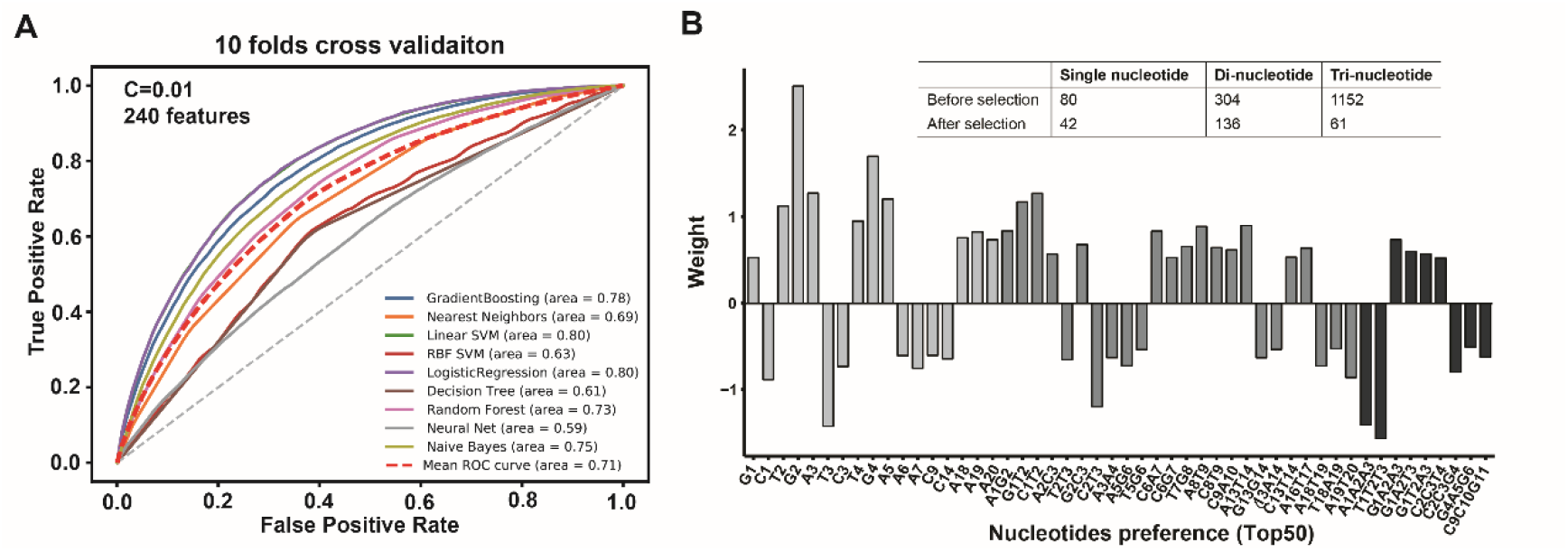
ROC curves of different models assessed and feature importance. **(A)** Performance in 9 models including gradient boosting, nearest neighboring, linear SVM, RBF SVM, logistic regression, decision tree, random forest, neural net and naïve bayes assessed by 10-folds cross validation. After feature selection by L1-SVM, only 240 features are incorporated in model selection. **(B)** Top 50 important features are depicted as bar plot. Weight of different features are shown in y-axis and different color means different type of features (single nucleotide, di-nucleotides and tri-nucleotides). The number of features before and after selection are also shown in upper panel.

### Webserver

We next created a webserver to package these two models for sgRNA selection as a web tool: OPT-sgRNA, a user-friendly website for sgRNA selection (Figure 5, http://bigdata.ibp.ac.cn/OPT-sgRNA/). It composes of sgRNA searching and pre-constructed libraries downloading. The web portal of OPT-sgRNA accepts more than one gene symbol or sequence as FASTA format as input, it helps user to design self-library with selected genes. Users can also set the desired numbers of off-target site to be evaluated and output of candidate sgRNAs, and select sgRNAs either Homo sapiens or Mus musculus. The background gene sequences are based on genome assembly hg38 (for human) and mm10 (for mouse), while pre-constructed libraries covering about 50,000 genes with 10 sgRNAs for each gene both for human and mouse are ready to be downloaded. The web portal is developed using in HTML and CSS scripts and implemented in Python based on the Django web framework, all of the backend scripts are written in Python programming language as well.

**Figure 5.**
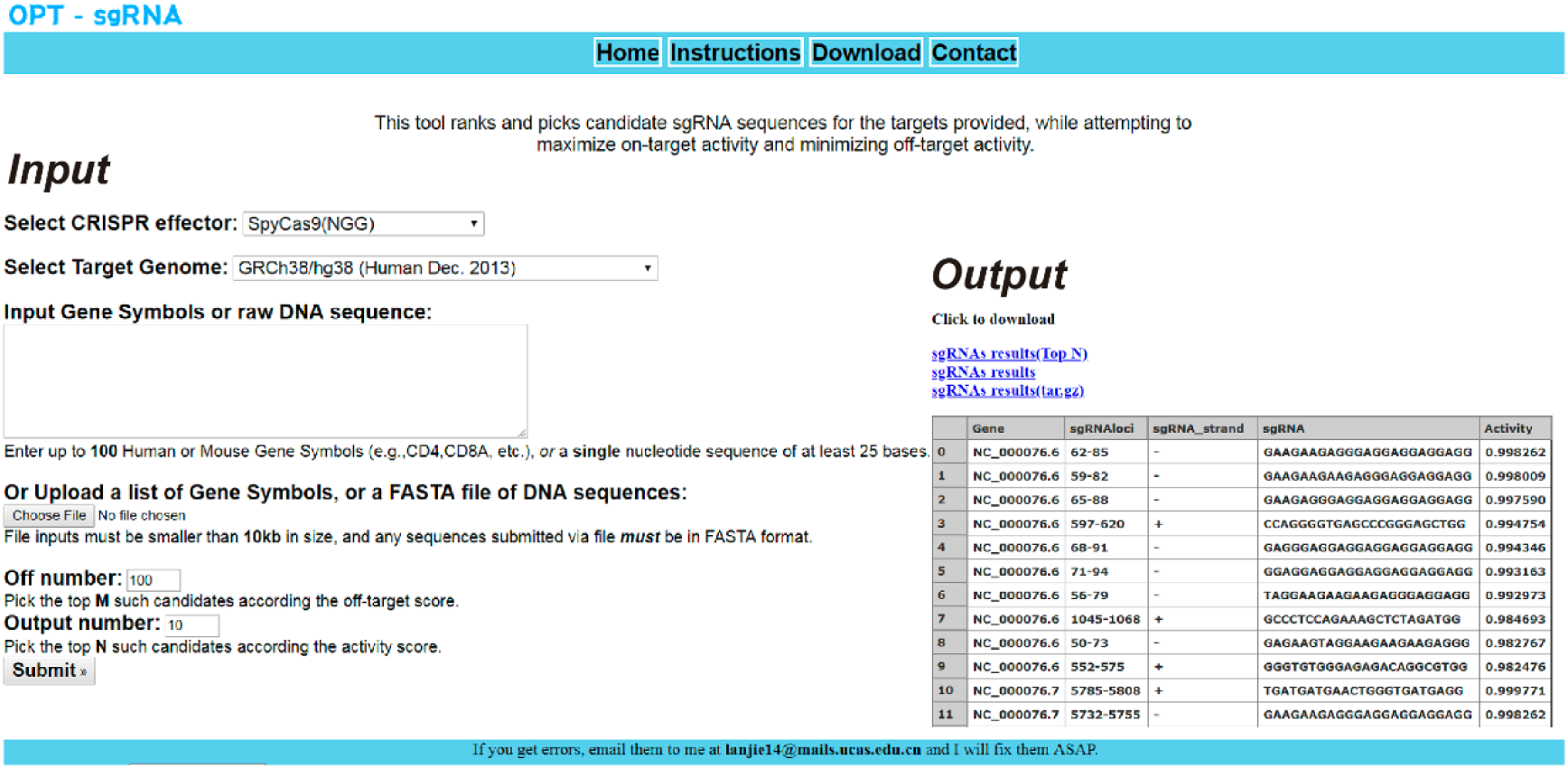
The main user interface of OPT-sgRNA website version. The input form for the multiple genes or sequences input consists of the parameters (reference genome, number of sgRNAs to be further scoring and number of output sgRNAs) for sgRNA selection and an example output was provided. Text file of gene list or multiple sequences also can also be uploaded as input.

## DISCUSSION

A successful sgRNA in any CRISPR/Cas9 genome editing experiments must maximize on-target activity and minimizing potential off-target effects at the same time. Therefore, computational tools for the prediction of on- and off-targeting assessment of sgRNAs is a crucial step for the efficient design of experiments. Although sevral tools have been developed for the design of sgRNAs and prediction of their off-target sites, they still could not robustly predict experimental success as prediction accuracy depends on the approximations of the underlying model and how closely the experimental setup matches the data the model was trained on.

In this work, we described a new open-source computational tool named OPT-sgRNA to design efficient sgRNAs with optimized target efficiency and specificity. There are several off-target prediction algorithms including CFD score, MIT score, CROP-IT score, CCTop score and Evaluation score. All of them are based on mismatches between sgRNA and target DNAs, and higher scores mean this sequence are more likely to be a true target site. Evaluation score is a penalized linear regression model based approach, and focus on mismatches (position and identity) and also including chromatin accessibility features. CCTop and CROP-IT are based on the distances of the mismatches to the PAM and CROP-IT incorporate penalty score for consecutive mismatches. We incorporated a new feature in the off-target effect evaluation: position weight of the mismatch sites. We found that 3-8 nt adjacent to PAM are the most important sits to despair sgRNAs activity. For the on-target activity scoring, we used random forest algorithm for feature selection. After feature filteration, only 240 features are incorporated in our model selection. And we observed several tri-nucleotides preference in in the middle of the sgRNA. Our optimized sgRNA design model provides a new method for the optimal design of efficient sgRNA and may improve the accuracy of sgRNA prediction.

## ACKNOWLEDGEMENT

We thank Dr. Yanli Wang for critical reading and helpful comments on the manuscript.

## REFERENCES

1. Jinek, M., Chylinski, K., Fonfara, I., Hauer, M., Doudna, J.A. and Charpentier, E. (2012) A programmable dual-RNA-guided DNA endonuclease in adaptive bacterial immunity. Science, 337, 816–821.

2. Cong, L., Ran, F.A., Cox, D., Lin, S., Barretto, R., Habib, N., Hsu, P.D., Wu, X., Jiang, W., Marraffini, L.A. et al. (2013) Multiplex genome engineering using CRISPR/Cas systems. Science, 339, 819–823.

3. Gilbert, L.A., Larson, M.H., Morsut, L., Liu, Z., Brar, G.A., Torres, S.E., Stern-Ginossar, N., Brandman, O., Whitehead, E.H., Doudna, J.A. et al. (2013) CRISPR-mediated modular RNA-guided regulation of transcription in eukaryotes. Cell, 154, 442–451.

4. Doudna, J.A. and Charpentier, E. (2014) Genome editing. The new frontier of genome engineering with CRISPR-Cas9. Science, 346, 1258096.

5. Hsu, P.D., Lander, E.S. and Zhang, F. (2014) Development and applications of CRISPR-Cas9 for genome engineering. Cell, 157, 1262–1278.

6. Wang, T., Wei, J.J., Sabatini, D.M. and Lander, E.S. (2014) Genetic Screens in Human Cells Using the CRISPR-Cas9 System. Science, 343, 80–84.

7. Wright, A.V., Nunez, J.K. and Doudna, J.A. (2016) Biology and Applications of CRISPR Systems: Harnessing Nature’s Toolbox for Genome Engineering. Cell, 164, 29–44.

8. Jinek, M., Jiang, F., Taylor, D.W., Sternberg, S.H., Kaya, E., Ma, E., Anders, C., Hauer, M., Zhou, K., Lin, S. et al. (2014) Structures of Cas9 endonucleases reveal RNA-mediated conformational activation. Science, 343, 1247997.

9. Nishimasu, H., Ran, F.A., Hsu, P.D., Konermann, S., Shehata, S.I., Dohmae, N., Ishitani, R., Zhang, F. and Nureki, O. (2014) Crystal structure of Cas9 in complex with guide RNA and target DNA. Cell, 156, 935–949.

10. Jiang, F., Zhou, K., Ma, L., Gressel, S. and Doudna, J.A. (2015) STRUCTURAL BIOLOGY. A Cas9-guide RNA complex preorganized for target DNA recognition. Science, 348, 1477–1481.

11. Sternberg, S.H., LaFrance, B., Kaplan, M. and Doudna, J.A. (2015) Conformational control of DNA target cleavage by CRISPR-Cas9. Nature, 527, 110–113.

12. Jiang, F., Taylor, D.W., Chen, J.S., Kornfeld, J.E., Zhou, K., Thompson, A.J., Nogales, E. and Doudna, J.A. (2016) Structures of a CRISPR-Cas9 R-loop complex primed for DNA cleavage. Science, 351, 867–871.

13. Yamano, T., Nishimasu, H., Zetsche, B., Hirano, H., Slaymaker, L.M., Li, Y., Fedorova, I., Nakane, T., Makarova, K.S., Koonin, E.V. et al. (2016) Crystal Structure of Cpf1 in Complex with Guide RNA and Target DNA. Cell, 165, 949–962.

14. Yamano, T., Zetsche, B., Ishitani, R., Zhang, F., Nishimasu, H. and Nureki, O. (2017) Structural Basis for the Canonical and Non-canonical PAM Recognition by CRISPR- Cpf1. Mol. Cell, 67, 633–645.

15. Zeng, Y., Cui, Y., Zhang, Y., Zhang, Y., Liang, M., Chen, H., Lan, J., Song, G. and Lou, J. (2018) The initiation, propagation and dynamics of CRISPR-SpyCas9 R-loop complex. Nucleic Acids Res., 46, 350–361.

16. Cui, Y., Tang, Y., Liang, M., Ji, Q., Zeng, Y., Chen, H., Lan, J., Jin, P., Wang, L., Song, G. et al. (2020) Direct observation of the formation of a CRISPR-Cas12a R-loop complex at the single-molecule level. Chem. Commun., 56, 2123–2126.

17. Fu, Y., Foden, J.A., Khayter, C., Maeder, M.L., Reyon, D., Joung, J.K. and Sander, J.D. (2013) High-frequency off-target mutagenesis induced by CRISPR-Cas nucleases in human cells. Nat. Biotechnol., 31, 822–826.

18. Hsu, P.D., Scott, D.A., Weinstein, J.A., Ran, F.A., Konermann, S., Agarwala, V., Li, Y., Fine, E.J., Wu, X., Shalem, O. et al. (2013) DNA targeting specificity of RNA-guided Cas9 nucleases. Nat. Biotechnol., 31, 827–832.

19. Pattanayak, V., Lin, S., Guilinger, J.P., Ma, E., Doudna, J.A. and Liu, D.R. (2013) High-throughput profiling of off-target DNA cleavage reveals RNA-programmed Cas9 nuclease specificity. Nat. Biotechnol., 31, 839–843.

20. Hendel, A., Fine, E.J., Bao, G. and Porteus, M.H. (2015) Quantifying on- and off-target genome editing. Trends Biotechnol., 33, 132–140.

21. Bradford, J. and Perrin, D. (2019) A benchmark of computational CRISPR-Cas9 guide design methods. PLoS Comput. Biol., 15, e1007274.

22. Wilson, L.O.W., O’Brien, A.R. and Bauer, D.C. (2018) The Current State and Future of CRISPR-Cas9 gRNA Design Tools. Front Pharmacol, 9, 749

23. Tycko, J., Myer, V.E. and Hsu, P.D. (2016) Methods for Optimizing CRISPR-Cas9 Genome Editing Specificity. Mol. Cell, 63, 355–370.

24. Liu, G., Zhang, Y. and Zhang, T. (2020) Computational approaches for effective CRISPR guide RNA design and evaluation. Comput. Struct. Biotechnol. J., 18, 35–44.

25. Peng, H., Zheng, Y., Blumenstein, M., Tao, D.C. and Li, J.Y. (2018) CRISPR/Cas9 cleavage efficiency regression through boosting algorithms and Markov sequence profiling. Bioinformatics, 34, 3069–3077.

26. Rahman, M.K. and Rahman, M.S. (2017) CRISPRpred: A flexible and efficient tool for sgRNAs on-target activity prediction in CRISPR/Cas9 systems. PLoS One, 12, e0181943.

27. Moreno-Mateos, M.A., Vejnar, C.E., Beaudoin, J.D., Fernandez, J.P., Mis, E.K., Khokha, M.K. and Giraldez, A.J. (2015) CRISPRscan: designing highly efficient sgRNAs for CRISPR-Cas9 targeting in vivo. Nat. Methods, 12, 982–988.

28. Biswas, A., Gagnon, J.N., Brouns, S.J., Fineran, P.C. and Brown, C.M. (2013) CRISPRTarget: bioinformatic prediction and analysis of crRNA targets. RNA Biol., 10, 817–827.

29. Kim, H.K., Min, S., Song, M., Jung, S., Choi, J.W., Kim, Y., Lee, S., Yoon, S. and Kim, H.H. (2018) Deep learning improves prediction of CRISPR-Cpf1 guide RNA activity. Nat. Biotechnol., 36, 239–241.

30. Heigwer, F., Kerr, G. and Boutros, M. (2014) E-CRISP: fast CRISPR target site identification. Nat. Methods, 11, 122–123.

31. Wilson, L.O.W., Reti, D., O’Brien, A.R., Dunne, R.A. and Bauer, D.C. (2018) High Activity Target-Site Identification Using Phenotypic Independent CRISPR-Cas9 Core Functionality. Crispr J., 1, 182–190.

32. Housden, B.E., Valvezan, A.J., Kelley, C., Sopko, R., Hu, Y., Roesel, C., Lin, S., Buckner, M., Tao, R., Yilmazel, B. et al. (2015) Identification of potential drug targets for tuberous sclerosis complex by synthetic screens combining CRISPR-based knockouts with RNAi. Sci. Signal., 8, rs9.

33. Doench, J.G., Fusi, N., Sullender, M., Hegde, M., Vaimberg, E.W., Donovan, K.F., Smith, I., Tothova, Z., Wilen, C., Orchard, R. et al. (2016) Optimized sgRNA design to maximize activity and minimize off-target effects of CRISPR-Cas9. Nat. Biotechnol., 34, 184–191.

34. Labuhn, M., Adams, F.F., Ng, M., Knoess, S., Schambach, A., Charpentier, E.M., Schwarzer, A., Mateo, J.L., Klusmann, J.H. and Heckl, D. (2018) Refined sgRNA efficacy prediction improves large- and small-scale CRISPR-Cas9 applications. Nucleic Acids Res., 46, 1375–1385.

35. Xu, H., Xiao, T., Chen, C.H., Li, W., Meyer, C.A., Wu, Q., Wu, D., Cong, L., Zhang, F., Liu, J.S. et al. (2015) Sequence determinants of improved CRISPR sgRNA design. Genome Res., 25, 1147–1157.

36. Chari, R., Yeo, N.C., Chavez, A. and Church, G.M. (2017) sgRNA Scorer 2.0: A Species-Independent Model To Predict CRISPR/Cas9 Activity. ACS Synth. Biol., 6, 902–904.

37. Zhang, D., Hurst, T., Duan, D. and Chen, S.J. (2019) Unified energetics analysis unravels SpCas9 cleavage activity for optimal gRNA design. Proc. Natl. Acad. Sci. U. S. A., 116, 8693–8698.

38. Chari, R., Mali, P., Moosburner, M. and Church, G.M. (2015) Unraveling CRISPR-Cas9 genome engineering parameters via a library-on-library approach. Nat Methods, 12, 823–826.

39. Wong, N., Liu, W. and Wang, X. (2015) WU-CRISPR: characteristics of functional guide RNAs for the CRISPR/Cas9 system. Genome Biol., 16, 218.

40. Singh, R., Kuscu, C., Quinlan, A., Qi, Y. and Adli, M. (2015) Cas9-chromatin binding information enables more accurate CRISPR off-target prediction. Nucleic Acids Res., 43, e118.

41. Bae, S., Park, J. and Kim, J.S. (2014) Cas-OFFinder: a fast and versatile algorithm that searches for potential off-target sites of Cas9 RNA-guided endonucleases. Bioinformatics, 30, 1473–1475.

42. Xiao, A., Cheng, Z., Kong, L., Zhu, Z., Lin, S., Gao, G. and Zhang, B. (2014) CasOT: a genome-wide Cas9/gRNA off-target searching tool. Bioinformatics, 30, 1180–1182.

43. Stemmer, M., Thumberger, T., Del Sol Keyer, M., Wittbrodt, J. and Mateo, J.L. (2015) CCTop: An Intuitive, Flexible and Reliable CRISPR/Cas9 Target Prediction Tool. PLoS One, 10, e0124633.

44. Jacquin, A.L.S., Odom, D.T. and Lukk, M. (2019) Crisflash: open-source software to generate CRISPR guide RNAs against genomes annotated with individual variation. Bioinformatics, 35, 3146–3147.

45. Alkan, F., Wenzel, A., Anthon, C., Havgaard, J.H. and Gorodkin, J. (2018) CRISPR- Cas9 off-targeting assessment with nucleic acid duplex energy parameters. Genome Biol., 19, 177.

46. Chuai, G., Ma, H., Yan, J., Chen, M., Hong, N., Xue, D., Zhou, C., Zhu, C., Chen, K., Duan, B. et al. (2018) DeepCRISPR: optimized CRISPR guide RNA design by deep learning. Genome Biol., 19, 80.

47. McKenna, A. and Shendure, J. (2018) FlashFry: a fast and flexible tool for large-scale CRISPR target design. BMC Biol., 16, 74.

48. Abadi, S., Yan, W.X., Amar, D. and Mayrose, I. (2017) A machine learning approach for predicting CRISPR-Cas9 cleavage efficiencies and patterns underlying its mechanism of action. PLoS Comput. Biol., 13, e1005807.

49. Listgarten, J., Weinstein, M., Kleinstiver, B.P., Sousa, A.A., Joung, J.K., Crawford, J., Gao, K., Hoang, L., Elibol, M., Doench, J.G. et al. (2018) Prediction of off-target activities for the end-to-end design of CRISPR guide RNAs. Nat. Biomed. Eng., 2, 38–47.

50. Xie, S., Shen, B., Zhang, C., Huang, X. and Zhang, Y. (2014) sgRNAcas9: a software package for designing CRISPR sgRNA and evaluating potential off-target cleavage sites. PLoS One, 9, e100448.

51. Hwang, G.H., Park, J., Lim, K., Kim, S., Yu, J., Yu, E., Kim, S.T., Eils, R., Kim, J.S. and Bae, S. (2018) Web-based design and analysis tools for CRISPR base editing. BMC Bioinformatics, 19, 542.

52. Ren, X., Yang, Z., Xu, J., Sun, J., Mao, D., Hu, Y., Yang, S.J., Qiao, H.H., Wang, X., Hu, Q. et al. (2014) Enhanced specificity and efficiency of the CRISPR/Cas9 system with optimized sgRNA parameters in Drosophila. Cell Rep., 9, 1151–1162.

53. Shalem, O., Sanjana, N.E., Hartenian, E., Shi, X., Scott, D.A., Mikkelsen, T.S., Heckl, D., Ebert, B.L., Root, D.E., Doench, J.G. et al. (2014) Genome-scale CRISPR-Cas9 knockout screening in human cells. Science, 343, 84–87.

54. Liu, X., Homma, A., Sayadi, J., Yang, S., Ohashi, J. and Takumi, T. (2016) Sequence features associated with the cleavage efficiency of CRISPR/Cas9 system. Sci. Rep., 6, 19675.

55. Doench, J.G., Hartenian, E., Graham, D.B., Tothova, Z., Hegde, M., Smith, I., Sullender, M., Ebert, B.L., Xavier, R.J. and Root, D.E. (2014) Rational design of highly active sgRNAs for CRISPR-Cas9-mediated gene inactivation. Nat. Biotechnol., 32, 1262–1267.

56. Klein, M., Eslami-Mossallam, B., Arroyo, D.G. and Depken, M. (2018) Hybridization Kinetics Explains CRISPR-Cas Off-Targeting Rules. Cell Rep., 22, 1413–1423.

57. Tsai, S.Q., Topkar, V.V., Joung, J.K. and Aryee, M.J. (2016) Open-source guideseq software for analysis of GUIDE-seq data. Nat. Biotechnol., 34, 483.

58. Kleinstiver, B.P., Pattanayak, V., Prew, M.S., Tsai, S.Q., Nguyen, N.T., Zheng, Z.L. and Joung, J.K. (2016) High-fidelity CRISPR-Cas9 nucleases with no detectable genome-wide off-target effects. Nature, 529, 490–495.

59. Chen, J.S., Dagdas, Y.S., Kleinstiver, B.P., Welch, M.M., Sousa, A.A., Harrington, L.B., Sternberg, S.H., Joung, J.K., Yildiz, A. and Doudna, J.A. (2017) Enhanced proofreading governs CRISPR-Cas9 targeting accuracy. Nature, 550, 407–410.

60. Rauscher, B., Heigwer, F., Breinig, M., Winter, J. and Boutros, M. (2017) GenomeCRISPR - a database for high-throughput CRISPR/Cas9 screens. Nucleic Acids Res, 45, D679–D686.

61. Aguirre, A.J., Meyers, R.M., Weir, B.A., Vazquez, F., Zhang, C.Z., Ben-David, U., Cook, A., Ha, G., Harrington, W.F., Doshi, M.B. et al. (2016) Genomic Copy Number Dictates a Gene-Independent Cell Response to CRISPR/Cas9 Targeting. Cancer Discov., 6, 914–929.

62. Jiang, H. and Wong, W.H. (2008) SeqMap: mapping massive amount of oligonucleotides to the genome. Bioinformatics, 24, 2395–2396.

