## Supplemental figures for "Optimized sgRNA design by deep learning to balance the off-target effects and on-target activity of CRISPR/Cas9"

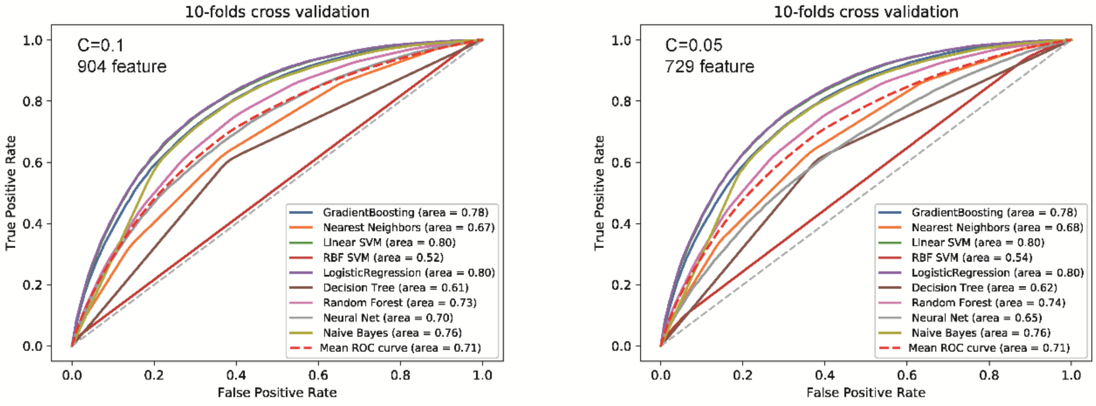


**Figure S1.** ROC curves of 9 models with different subset of features including gradient boosting, nearest neighboring, linear SVM, RBF SVM, logistic regression, decision tree, random forest, neural net and naïve bayes assessed by 10-folds cross validation.

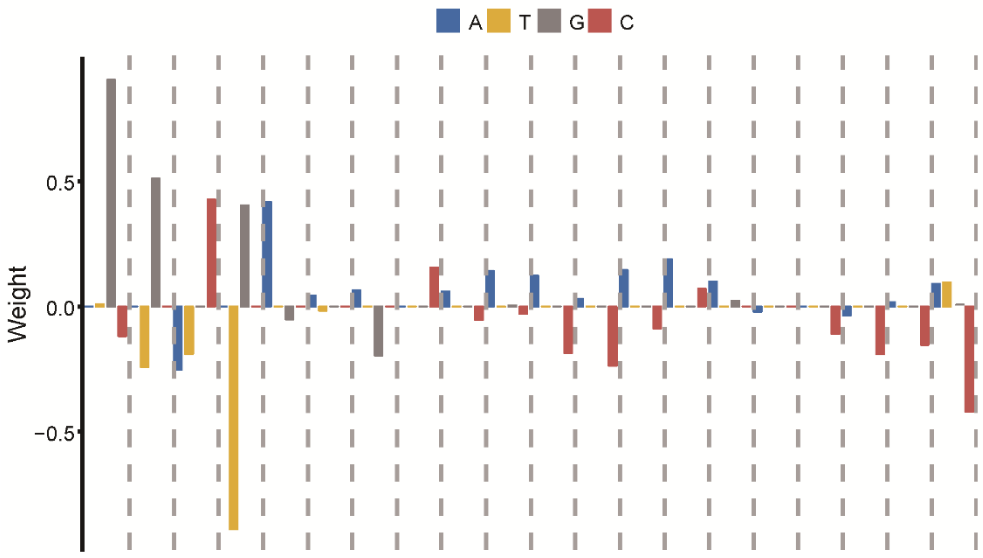


**Figure S2.** Nucleotide preference assessed by the logistic regression. The y-axis direction indicates whether a given nucleotide is favored or disfavored for cleavage activity.
